# Pathogenic LRRK2 regulates ciliation probability upstream of Tau Tubulin kinase 2

**DOI:** 10.1101/2020.04.07.029983

**Authors:** Yuriko Sobu, Paulina S. Wawro, Herschel S. Dhekne, Suzanne R. Pfeffer

## Abstract

Mutations that activate LRRK2 protein kinase cause Parkinson’s disease. We have shown previously that Rab10 phosphorylation by LRRK2 enhances its binding to RILPL1 and together, these proteins block cilia formation in a variety of cell types including patient derived iPS cells. We have used live cell fluorescence microscopy to identify, more precisely, the effect of LRRK2 kinase activity on both the formation of cilia triggered by serum starvation and loss of cilia seen upon serum re-addition. LRRK2 activity decreases the overall probability of ciliation without changing the rates of cilia formation in R1441C LRRK2 MEF cells. Cilia loss in these cells is accompanied by ciliary decapitation. Kinase activity does not change the timing or frequency of decapitation or the rate of cilia loss, but increases the percent of cilia that are lost upon serum addition. LRRK2 activity, or overexpression of RILPL1 protein, blocks release of CP110 from the mother centriole, a step normally required for early ciliogenesis. In both cases, failure of CP110 uncapping was due to failure to recruit TTBK2, a kinase needed for CP110 release. In contrast, recruitment of EHD1, another step important for ciliogenesis, appears unaltered. These experiments provide critical detail to our understanding of the cellular consequences of pathogenic LRRK2 mutation, and indicate that LRRK2 blocks ciliogenesis upstream of TTBK2 and enhances the deciliation process in response to serum addition.

**SIGNIFICANCE STATEMENT:** Mutations that activate LRRK2 protein kinase cause Parkinson’s disease. LRRK2 phosphorylates a subset of Rab GTPases, in particular Rab8 and Rab10. Phosphorylated Rabs bind preferentially to a distinct set of effectors and block in primary ciliation in multiple cell types. We show here that the cilia blockade is upstream of the recruitment of TTBK2 kinase to the mother centriole, a step required for the release of CP110 and subsequent cilia formation. This study provides fundamental information related to how pathogenic LRRK2 interferes with normal cell physiology.

## >Introduction

Activating mutations in LRRK2 kinase are responsible for inherited Parkinson’s disease. Pathogenic LRRK2 phosphorylates a subset of Rab GTPases (1, 2) that are master regulators of protein transport through the secretory and endocytic pathways (3, 4). Upon phosphorylation, these Rab GTPases bind to a new set of effector proteins (1) triggering likely diverse physiological events.

Together with the labs of Dario Alessi and Matthias Mann, we have shown that LRRK2 kinase activity blocks primary cilia formation in cultured cells (1) and in specific areas of mouse brains carrying pathogenic LRRK2 mutations (5). Inhibition of cilia formation requires Rab10 and a protein, RILPL1 (5) that binds with strong preference to phosphorylated Rab10 protein (1). This process is regulated by PPM1H phosphatase that is highly specific for pRab8A and pRab10 proteins (6). LRRK2-mediated Rab phosphorylation of Rab proteins has also been reported to alter the association of mother and daughter centrioles (7). Rab phosphorylation appears to be part of the normal process by which cilia formation is regulated, both in wild type cells and even more so, in cells expressing hyperactive mutant LRRK2 kinase, as depletion of PPM1H phosphatase is sufficient to block cilia formation (6).

Cilia growth and resorption are closely linked to the cell cycle. Upon serum starvation, removal of CP110 from the mother centriole is a key step in the initiation of cilia formation (8). Tau tubulin kinase 2 (TTBK2), a protein mutated in the inherited movement disorder, Spinocerebellar ataxia type 11, promotes the removal of CP110 and recruitment of IFT proteins to build the ciliary axoneme (8–10). TTBK2 is recruited selectively to the mother centriole by binding to CEP164 (11, 12). Huang et al. (13) showed that TTBK2 triggers ciliogenesis by phosphorylating MPP9, leading to MPP9 degradation and subsequent release of CP110/CEP97. TTBK2 also binds and phosphorylates CEP83 (14), further facilitating CP110 release and cilia formation.

We have carried out live cell microscopy experiments to investigate the mechanisms by which LRRK2 activity inhibits cilia formation in cell culture and find that the blockade is early, upstream of TTBK2 recruitment to the mother centriole.

## Results

To better characterize the step(s) of cilia formation or disassembly altered by LRRK2 mediated Rab10 phosphorylation, we carried out live cell video microscopy of knock-in MEF cells expressing pathogenic R1441C LRRK2 at endogenous levels; these cells were engineered to stably express the cilia-localized G protein coupled receptor, SSTR3-GFP (15) and a pericentrin-AKAP-450 centrosomal targeting domain (PACT, 16) tagged with mKO2 (red in Fig. 1A). To monitor cilia formation, cells were transferred to serum free medium and filmed for 8 hours, with or without a LRRK2 specific inhibitor, MLi-2, with images captured every 15 minutes. We tracked cilia using SSTR3 instead of the common marker, Arl13B since exogenous expression of Arl13b caused elongated primary cilia and altered their stability.

**Fig. 1.**
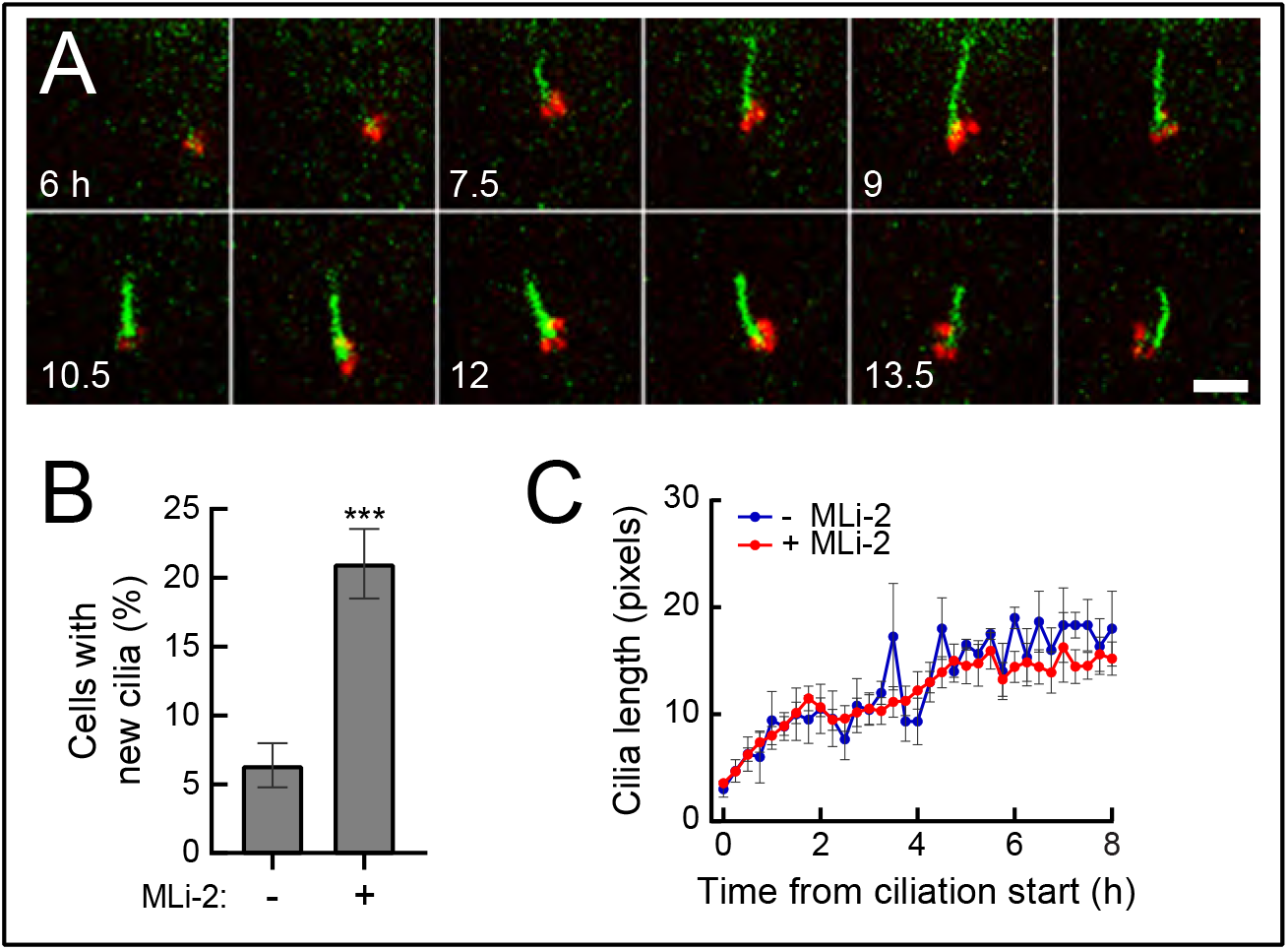
Pathogenic LRRK2 activity decreases the probability of cilia formation. **A.** Time-lapse images of MEF R1441C cells stably expressing SSTR3-GFP (green) and mKO2-PACT (red). After 2hr of serum starvation ± MLi-2, cells were imaged every 15 min. In this and subsequent figures, intermediate-time images were removed from the figure to conserve space. Scale bar, 5μm. **B.** Percentage of cells that generated primary cilia ± MLi-2 over 8hr of imaging. Values represent the mean ±SEM from 3 independent experiments, each with > 20 cells. Significance was determined by t-test; ***, P = 0.0026 **C.** Rate of cilia growth. Lengths of cilia were measured after SSTR3^+^ cilia appeared (t=0). Values represent the mean ± SEM from 3-7 (-MLi-2) and 9-21 (+MLi-2) cilia (cilia numbers were different at different time points over 8 hr).

Addition of the LRRK2 inhibitor, MLi-2, significantly increased the number of cells that generated a cilium under these low confluency conditions (22% versus 7%), as expected (Fig. 1A,B). Focusing only on cells that grew cilia, the rate of cilia growth was indistinguishable in the presence or absence of LRRK2 inhibitor, plateauing after about 3 hours of serum starvation (Figure 1C). If a cell ciliated, the rate of growth was the same, independent of LRRK2 activity. Thus, pathogenic LRRK2 influences the probability that a cell will initiate cilia growth upon serum starvation.

To monitor cilia loss from LRRK2 mutant cells, cilia formation was first initiated by serum starvation (~24h) and then cells were transferred to serum containing medium ± MLi-2 (Fig. 2A, F). This protocol enabled us to monitor the properties of the smaller proportion of cilia formed despite the presence of mutant LRRK2 activity; the process by which these cilia were lost could then be followed with or without active kinase. Unlike RPE cells, deciliation in MEFs includes “ectocytosis” or decapitation from the tips of cilia (17–19); yellow arrowheads in Fig. 2A).

**Fig. 2.**
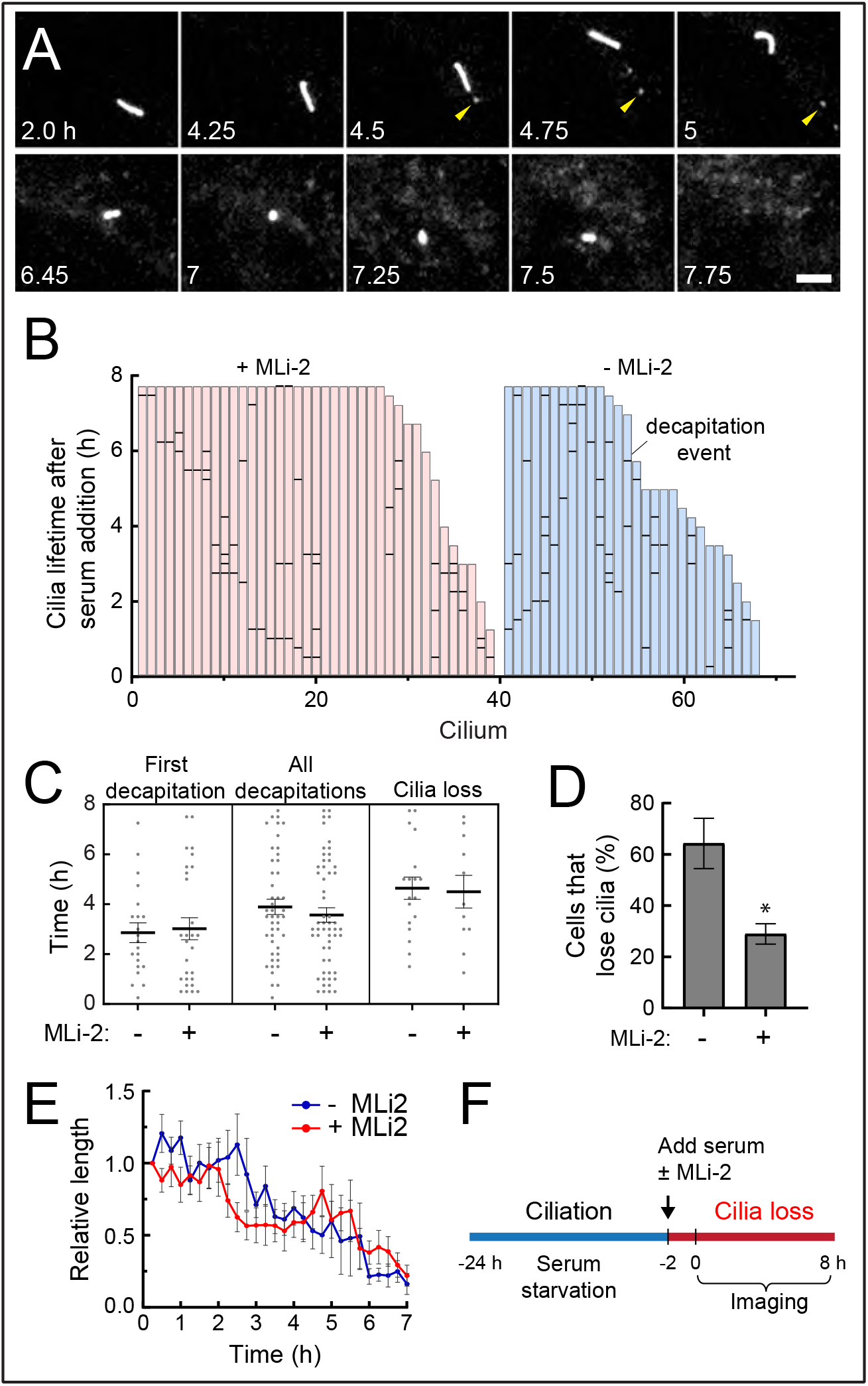
Cilia loss upon serum re-addition is enhanced by pathogenic LRRK2 activity. **A.** Time-lapse images of R1441C LRRK2 MEF cells stably expressing SSTR3-GFP. Cells were serum starved for 24hr and cilia loss was triggered by serum re-addition ± MLi-2. After 2hr, cells were imaged every 15 minutes. Arrowheads indicate decapitated vesicles. Scale bar, 5μm. **B.** Individual bars indicate single cilia. Heights of the bars (red, +MLi-2, Blue, -MLi-2) represent the time at which a given cilium disappeared. Horizontal black bars indicate decapitation events. 27 MLi-2^+^ cilia and 11 MLi-2^−^ cilia remained; 11 MLi-2^+^ cilia and 18 MLi-2^−^ cilia disappeared during the imaging period. **C.** Average timing of events. Times of first decapitation for each cilium (P = 0.80), all decapitations (P = 0.45), and cilia loss (P = 0.85) are shown. Values represent the mean ± SEM. Significance was determined by t-test. **D.** Probability of cilia loss during imaging is shown. Values represent the mean ± SEM from 3 independent experiments each containing > 7 cells. Significance was determined by t-test; *, P = 0.029. **E.** Rates of cilia loss. Lengths of cilia that disappeared during imaging were measured. Values represent the mean ± SEM from 3 to 18 (-MLi-2) and 3 to 11 (+MLi-2) cilia. **F.** Timeline of this imaging experiment.

Each bar in Fig. 2B represents the lifetime of an individual cilium, with horizontal lines indicating the timing of decapitation events. The left-most bars in each set (red or blue) indicate cilia that were stable over the imaging time frame; the adjacent bars of decreasing height represent cilia with the indicated, shorter lifetimes (<8 hours). Note that in the absence of MLi-2 (blue bars), there were fewer stable cilia and proportionately more cilia loss. As quantified in Fig. 2C, we detected no significant difference in terms of the times at which a cilium displayed its first decapitation event ± MLi-2. In addition, the total number of subsequent decapitation events and their timing were similar, with or without LRRK2 activity. Thus, LRRK2 does not appear to influence the overall timing or pathway of decapitation. Nevertheless, LRRK2 activity increased the proportion of cells that lost their cilia over an 8 hour observation window (Figs. 2B, D). For those cells that lost cilia, the rate of loss was similar ± MLi-2 (Fig. 2E; >9 cells averaged for each curve). What differed was the overall probability of deciliation within the first 8 hours of serum addition, which was 64% without inhibitor and 29% with inhibitor (Fig. 2D). LRRK2 activity led to a larger proportion of cells losing their cilia, as if kinase activity facilitated events that trigger cilia loss upon serum addition.

We next examined the deciliation process for cilia that were formed without kinase activity (+MLi-2) during serum starvation, followed by observation with or without LRRK2 inhibition in the continued absence of serum (Fig 3). This protocol examines the intrinsic stability of cilia after LRRK2 reactivation by MLi-2 washout. LRRK2 activity had no influence on the percent of cells with persistent cilia (Fig. 3B), the percent of cells displaying decapitation (Fig 3C), the stability of cilia length (Fig 3D), the time of first decapitation or the overall frequency of decapitations (Fig 3E). These experiments confirm that the loss of cilia seen in the presence of pathogenic LRRK2 activity relates to the process by which serum addition initiates cilia loss.

**Fig. 3.**
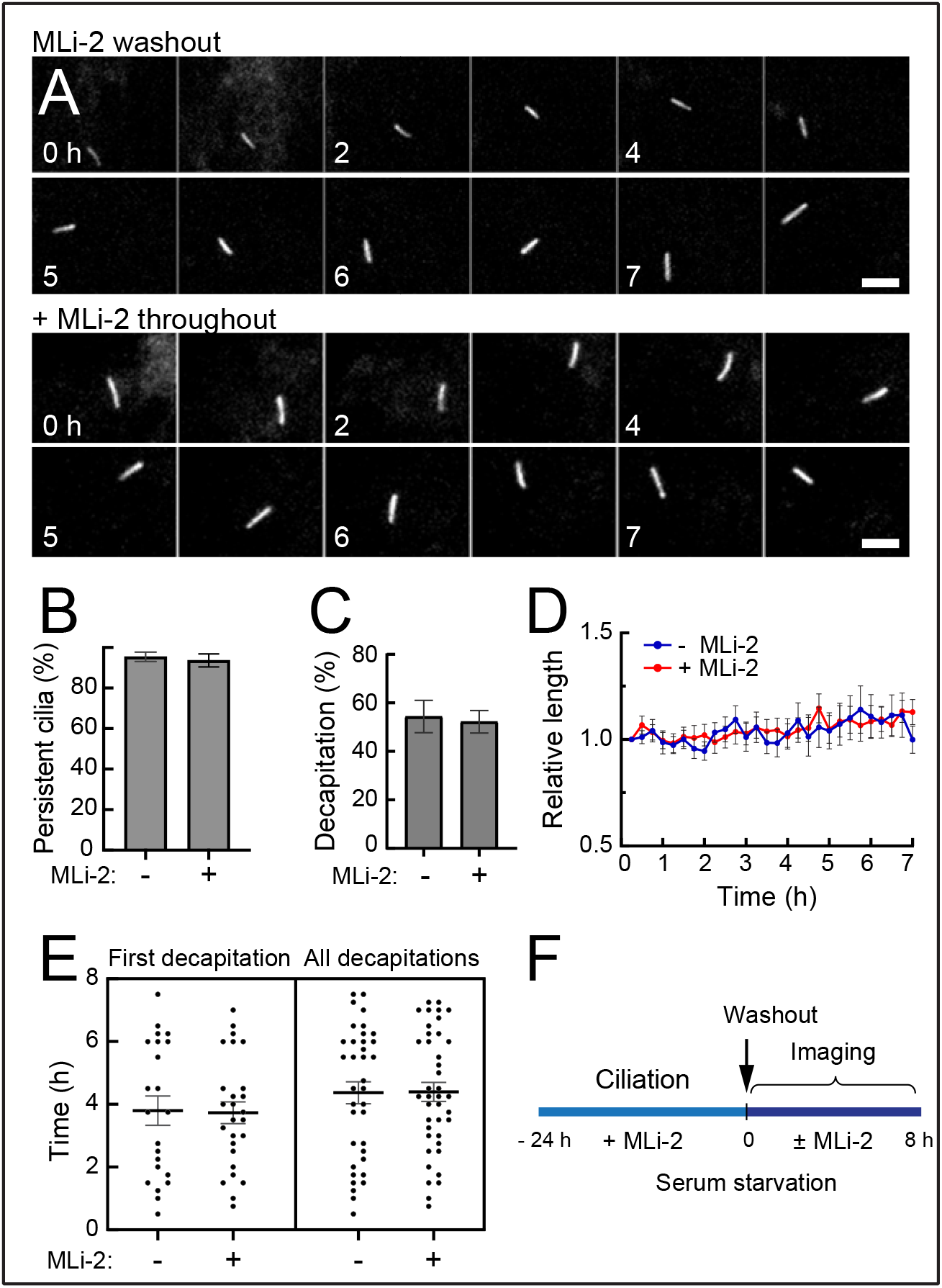
Pathogenic LRRK2 activity does not alter cilia stability under serum starvation conditions. **A.** Time-lapse images of R1441C LRRK2 MEF cells stably expressing SSTR3-GFP. Cells were serum starved in the presence of MLi-2 for 24hr to generate “wild type” cilia. After drug washout, cells were imaged every 15 minutes under continued serum starvation ± MLi-2. Scale bar, 5μm. Times are indicated in hours. **B**. Probability of persistent cilia (P=0.68) and **C**. Percentage of cilia that displayed decapitation events (P=0.80) during 8 hr of imaging are shown. Values represent the mean ± SEM from 3 independent experiments, each containing >10 cells. Significance was determined by t-test. **D.** Relative growth of cilia after MLi-2 washout. Values represent the mean ± SEM from 39 cells +MLi-2 and 50 cells -MLi-2. **E.** Timing of first decapitation of each cilium (P=0.91) and all decapitations (P=0.96) are shown. Values represent the mean ± SEM. Significance was determined by the t-test (NS). **F.** Timeline of this imaging experiment.

Similar findings were obtained if cilia from R1441C LRRK2 MEFs were generated with or without LRRK2 activity in the continued absence of serum (Supplemental Figure 1). Cilia that formed despite the presence of pathogenic kinase were just as persistent as those formed in the presence of kinase inhibitor, over a 8 hour observation period (Supp Fig. 1C). Again, cilia showed comparable timing of first and total decapitation events and a similar fraction of total cilia displayed decapitation. Altogether, these data show that pathogenic LRRK2 influences cilia formation propensity and sensitivity to serum triggering the cilia resorption and loss process.

### Mechanism of ciliation blockade

Removal of CP110 from the mother centriole is a key step in the initiation of cilia formation (8, 20). CP110 is recruited to the distal end of the centriole by binding to CEP97; the CP110-CEP97 complex is recruited to centrioles by MPP9 and together they suppress microtubule assembly (13). We therefore explored CP110 status at the mother centriole to check its relationship to the decreased probability of ciliogenesis in cells expressing pathogenic LRRK2.

Fig. 4 shows that in the presence of pathogenic LRRK2, upon overnight serum starvation conditions that reveal a ciliogenesis defect, both mother and daughter centrioles were capped with CP110 protein in ~75% of cells. Treatment with MLi-2 to inhibit LRRK2 kinase reversed this block, enabling CP110 release from the mother centriole and subsequent ciliogenesis in 77% of cells, within 24 hours of treatment.

**Fig. 4.**
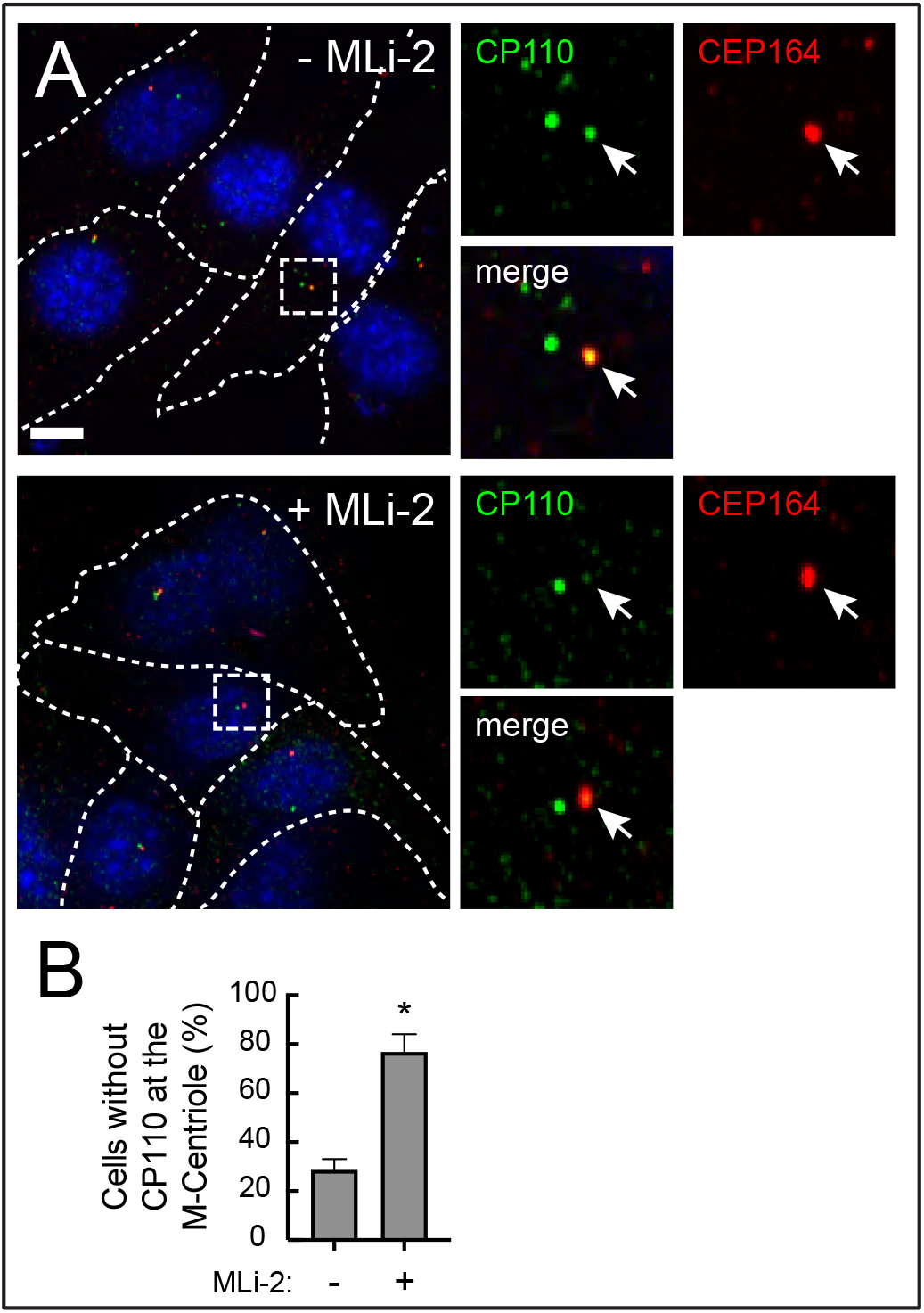
Pathogenic LRRK2 activity blocks release of CP110 from the mother centriole. **A.** R1441C LRRK2 MEF cells were starved for 24h ± MLi-2. Cells were then fixed with cold methanol and stained with rabbit anti-CP110 (green) and mouse anti-CEP164 (red) antibodies. Smaller images at right are enlarged regions boxed in the larger images at left. **B.** Percent of cells lacking a CP110 cap at the mother centriole, determined by CEP164 staining. Values represent the mean and SEM of two independent experiments (n > 30 in each experiment). Significance was determined by t-test. *, P = 0.0295. Scale bars, 10μm.

To test if pathogenic LRRK2 influenced the recruitment of TTBK2 to the mother centriole, we stably introduced GFP tagged TTBK2 at barely detectable levels into R1441C LRRK2 MEF cells and monitored its recruitment to the mother centriole upon serum starvation. (Endogenous TTBK2 was not detectable by immunofluorescence staining or western blot using a commercial antibody in this cell type). In addition to R1441C LRRK2 expression blocking release of CP110 from mother centrioles in MEF cells (Fig. 4), hyperactive LRRK2 also blocked the recruitment of TTBK2 to the centriolar region marked by gamma-tubulin (Fig. 5) and more specifically, to the mother centriole, marked by distal appendage protein CEP164 (Fig. 6). In contrast, in cells treated overnight with the LRRK2 specific MLi-2 inhibitor, 90% of cells showed a bright TTBK2 spot at the mother centriole, as expected for cells in which CP110 is released.

**Fig. 5.**
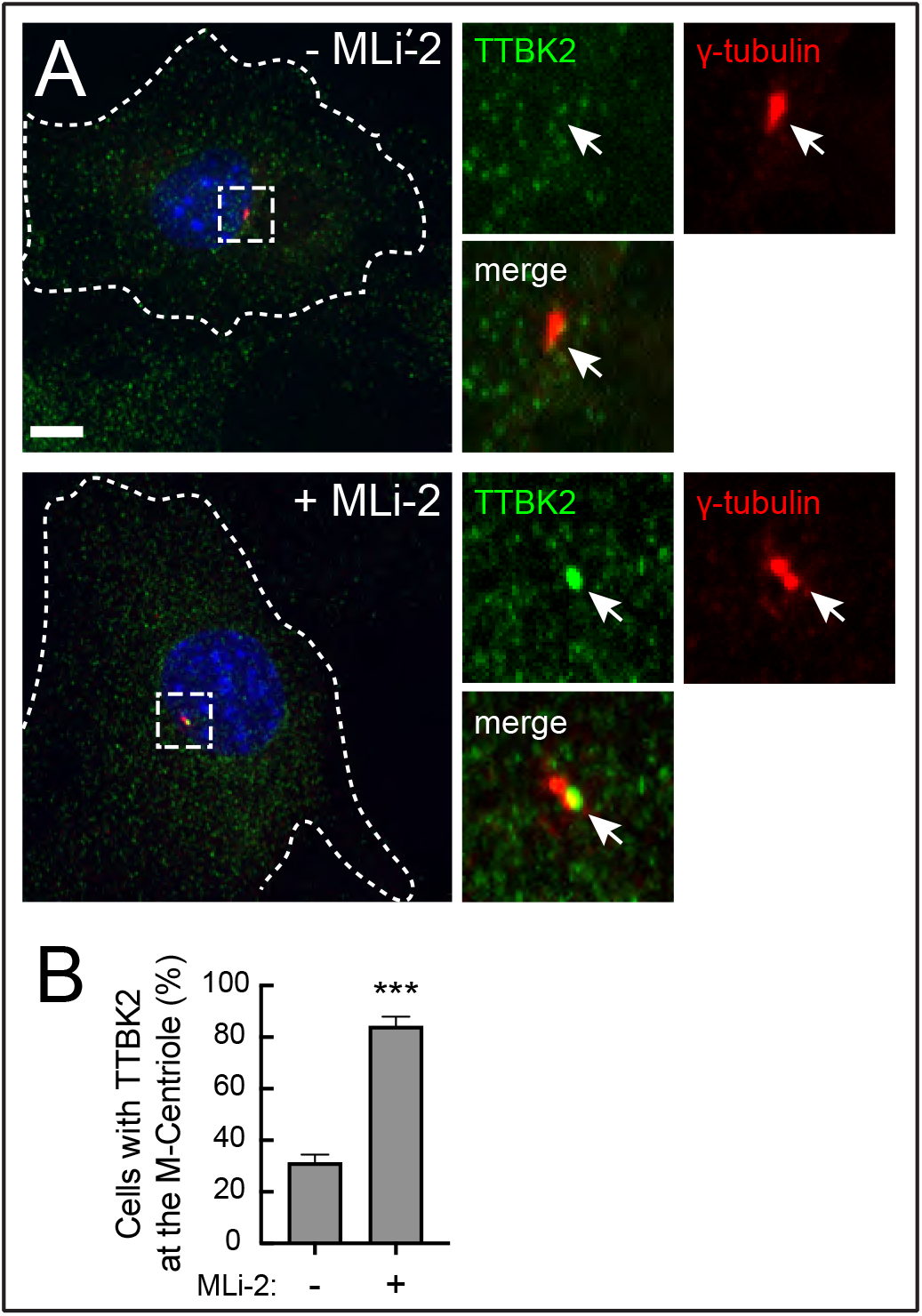
Pathogenic LRRK2 activity blocks TTBK2 recruitment to the mother centriole. **A.** R1441C LRRK2 MEF cells stably expressing GFP-TTBK2 were starved for 24h ± MLi-2. Cells were then fixed with cold methanol and stained with rabbit anti-TTBK2 (green) and mouse anti-γ-tubulin (red) antibodies. At right are enlarged regions boxed in the larger images at left. **B.** Percent of cells with TTBK2 at the centrosome, marked by γ-tubulin staining. Values represent the mean and SEM of two independent experiments (n>60 in each experiment). Significance was determined by the t-test; ***, P = 0.0002. Scale bars, 10μm.

**Fig. 6.**
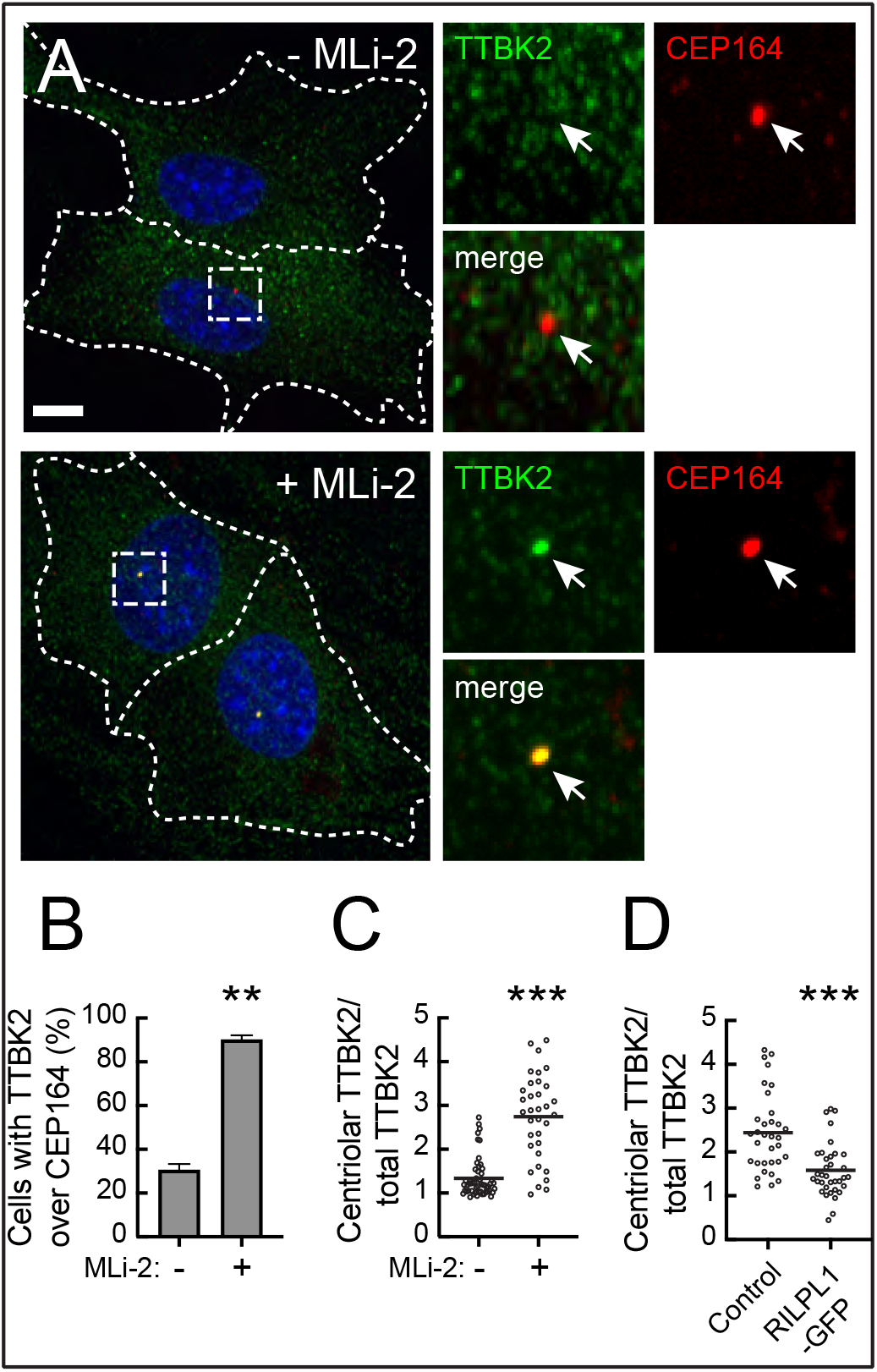
Pathogenic LRRK2 activity blocks TTBK2 recruitment without affecting CEP164. **A.** R1441C LRRK2 MEF cells stably expressing GFP-TTBK2 were starved for 24h with 200nM MLi-2 or DMSO. Cells were then fixed with cold methanol and stained with rabbit anti-TTBK2 (green) and mouse anti-CEP164 (red) antibodies. At right are enlarged regions boxed in the larger images at left. **B.** Percent of cells with TTBK2 at the mother centriole, marked by CEP164 staining. Values represent the mean and SEM of two independent experiments (n > 30 in each experiment). Significance was determined by the t-test; **, P = 0.003. **C.** Each dot represents the ratio of mean TTBK2 signal intensity at the centriolar area divided by the mean signal intensity throughout the cell (n > 30). Horizontal lines represent mean. P = 0.0001. **D.** RPE cells were transfected with RILPL1-GFP; its expression was induced with 1μg/ml doxycycline and cells were serum starved for 24h, methanol fixed and stained for endogenous TTBK2 and CEP164. Cells were then imaged and analyzed as in C. Each dot represents the ratio of mean TTBK2 signal intensity at the centriolar area divided by the mean signal intensity throughout the cell (n > 40). Significance was determined by the t-test, P = 0.0001. Scale bars, 10μm.

In untreated R1441C LRRK2 MEF cells, we often saw a weak signal of TTBK2 adjacent to gamma-tubulin positive centrioles which we ascribe to the established localization of TTBK2 to the plus-ends of microtubules (21,22). Lo et al. (14) also report a scattered radial signal of TTBK2 around the mother centriole in cells without serum starvation. We quantified the enrichment of TTBK2 over mother centriole (marked by CEP164) using CellProfiler. Using an unbiased image analysis pipeline, we detected an average 2-fold enrichment of TTBK2 over the mother centriole upon MLi-2 treatment (Fig. 6C; same conditions as in Figs. 5A, 6A).

We have shown previously that LRRK2 phosphorylated Rab10 blocks ciliation in concert with its binding partner, RILPL1; RILPL1 overexpression is also sufficient to dominantly block cilia formation in the absence of pathogenic LRRK2 (5). Since RILPL1 localizes adjacent to mother centriole, (5, 23) we tested whether RILPL1 overexpression can also influence TTBK2 recruitment. In RPE cells where (unlike MEFs) it was possible to quantify endogenous TTBK2 at the CEP164 marked mother centriole, expression of RILPL1-GFP reduced TTBK2 recruitment to the mother centriole two fold compared with control cells (Figure 6D and Supplemental Fig. 2). Thus, exogenous expression of RILPL1-GFP phenocopies the effect of pathogenic LRRK2 on centriolar TTBK2 recruitment.

Early membrane trafficking events that include recruitment of distal appendage vesicles (DAVs) and ciliary vesicles (CVs) are critical for axoneme extension (24). EHD1 is a membrane associated protein needed for both ciliary vesicle formation and release of CP110 from the distal end of mother centriole (25). Despite the retention of CP110 at the mother centriole in mutant LRRK2 expressing cells, we found that most cells retained EHD1 staining near centrosomes, and centrosomal EHD1 staining levels did not change upon LRRK2 inhibition (Fig. 7). These data suggest that the ability of LRRK2 to block CP110 release from the mother centriole is independent of EHD1 recruitment to the centrosomal region, and is more closely linked to failure of TTBK2 recruitment rather than to a block in the ciliary vesicle fusion pathway.

**Fig. 7.**
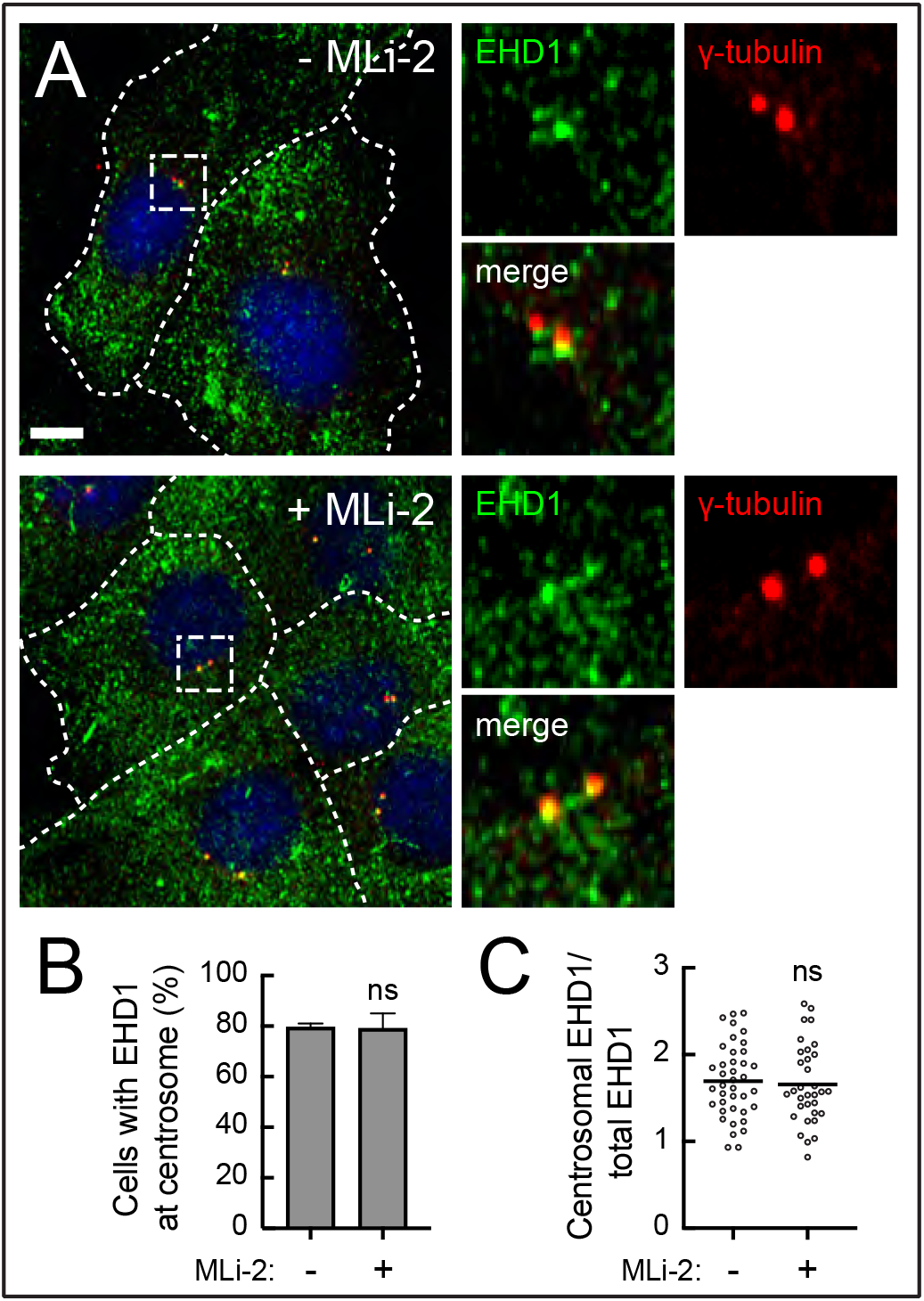
LRRK2 activity does not change EHD1 concentration at the centrosome. **A.** R1441C LRRK2 MEF cells stably expressing GFP-TTBK2 were starved for 24h ± MLi-2. Cells were then fixed with cold methanol and stained with rabbit anti-EHD1 (green) and mouse anti-γ-tubulin (red) antibodies. At right are enlarged regions boxed in the larger images at left. Scale bars, 10μm. **B.** Percent of cells with EHD1 at the centrosome, marked by γ-tubulin staining. Values represent the mean and SEM of two independent experiments (n > 30 in each experiment). Significance was determined by the t-test; P = 0.9429. **C.** Each dot represents the ratio of mean EHD1 signal intensity at the centrosome divided by the mean signal intensity throughout the cell (n > 40). Horizontal lines represent mean. P = 0.7225 (ns).

## Discussion

We have shown here that pathogenic LRRK2 decreases the probability that a cilium will form under conditions of serum starvation and the probability of cilia loss in mouse embryo fibroblasts. This is because TTBK2 fails to be recruited to the mother centriole in the presence of pathogenic LRRK2 activity. Under these conditions, CP110 continues to cap both mother and daughter centrioles, thereby blocking axoneme extension. When deciliation was examined by live cell microscopy, we found that R1441C LRRK2 MEF cells lost cilia by a process of retraction and ectocytosis or decapitation, independent of whether pathogenic LRRK2 was expressed. We saw no effect of LRRK2 activity on the timing of the first decapitation event or the number or timing of all subsequent decapitation events. Thus, LRRK2 does not influence the pathway of cilia loss. Rather, LRRK2 mutant-expressing cells seem more sensitive to serum-triggered initiation of deciliation and cell cycle progression.

CP110 is a negative regulator of cilia formation that functions with CEP97 to restrict the ciliogenesis program to the appropriate stage of the cell cycle (20). Multiple cellular components drive CP110 release from the mother centriole prior to microtubule extension, via TTBK2 recruitment and the formation of a ciliary vesicle. Although EHD1 is essential for ciliary vesicle-associated decapping (25), we saw no apparent change in EHD1 accumulation adjacent to the mother centriole. This suggests that the inability to recruit TTBK2 to the mother centriole and interact with CEP164 is the predominant pathway by which primary cilia formation is blocked in LRRK2 mutant MEF cells.

We had previously shown that LRRK2 generated, phospho-Rab10 GTPase and it’s novel binding partner, RILPL1, are both required for the process by which LRRK2 blocks cilia (5). RILPL1 is centriolar (23), as are complexes of pRab10 and RILPL1 protein (5). In addition, RILPL1 is localized in part to the centriole, even in cells lacking Rab10 protein (5). Indeed, we found that simple overexpression of RILP1 that phenocopies mutant LRRK2 expression (5) also decreases TTBK2 recruitment to the mother centriole. It is very likely that the pRab10-RILPL1 complex acts as a dominant negative inhibitor of TTBK2 recruitment, possible by occluding interaction of this kinase with CEP164.

The precise signals that lead to TTBK2 recruitment to the mother centriole and regulate its subsequent interaction with CEP164 are still unclear. Xu et al. (26) presented a model in which phosphatidylinositol 4-phosphate (PI4P) levels adjacent to the mother centriole regulate the interaction of TTBK2 with CEP164. INPP5E phosphatase generates PI4P from PI4,5P adjacent to the mother centriole; PIPKI gamma re-converts the PI4P to PI4,5P2. It is possible that LRRK2 phosphorylation changes the levels of phosphoinositide kinases and phosphatases present at the mother centriole to impact this interaction. In this regard, OCRL and INPP5B are both Rab8 effectors that may be lost upon Rab8 phosphorylation, as phosphorylation decreases the ability of Rab8 to bind either of these proteins (2). Future experiments will enable us to explore these various scenarios in deeper mechanistic detail.

## Materials and Methods

### Plasmids

DNA constructs were amplified in E. coli DH5a and purified using mini prep columns (Enzymax). DNA sequence verification of all plasmids was performed by Sequetech (http://www.sequetech.com). PACT-mKO2 was generated by amplifying the PACT domain of Pericentrin from cDNA of RPE cells using the primers cggatatccatcacactggcggccgcatggaccccggccggct and ttgatccctcgatgttaactctagattaatcatcgggtggcaggatttct and cloned into Not1/Xba1 linearized lentiviral vector CSII-EF-mKO2 (gift of Aaron Straight, Stanford) using Gibson assembly cloning. PACT domain was determined as described by Gillingham AK, EMBO Reports 2000. mApple-SSTR3-N-17 was a gift from Michael Davidson (Addgene # 54949). SSTR3 was shuttled into CSII-EF lentiviral vector into XhoI/XbaI site by Gibson cloning using the primers aacacgctaccggtctcgagaattcatggccactgttacctatcctt and tgctcaccatcagatggctcagtgtgctgg for SSTR3, gagccatctgatggtgagcaagggcgag and ttgatccctcgatgttaactctagattacttgtacagctcgtccatgcc for GFP sequence from the EGFP-N1 vector. eGFP-TTBK2 wild type (DU17481) and kinase dead mutant (DU17482) were obtained from MRC-Protein and phosphorylation unit, Dundee, Scotland and cloned into modified lentiviral vector pMCB306 into added restriction site AscI/EcoRI by Gibson assembly using primers cgagctgtacaaggggtccggactcagatccac and attaggtccctcgacgctatctgctgagtttactggctgg. RILPL1-GFP was cloned into pcDNA5D-TO and expression induced with Doxycycline (1, 5).

### Antibodies and reagents

Rabbit anti-TTBK2 (Sigma, HPA018113, 1:2000), mouse anti-Arl13b (Neuromab, 75-287, 1:2000), mouse anti-Gamma tubulin (Invitrogen, MA1-19421, 1:2000), Rabbit anti-CP110 (Proteintech, 12780-1-AP, 1:2000), mouse anti-CEP164 (Santa Cruz, sc-515403, 1:1000), rabbit anti-EHD1 (Abcam, ab109747, 1:1000), mouse anti-GFP (Neuromab, 73-131, 1:2000), mouse anti-mCherry (Novus, NBP1-96752, 1:1000), chicken anti-GFP (Aves, GFP1020, 1:5000). MLi-2 was obtained from Tocris Biosciences (Cat no 5756) and dissolved in DMSO to 1mM stock and kept in −20°C.

Cell culture, transfection and viral transduction –Mouse embryonic fibroblasts (MEFs) derived from LRRK2-R1441C knock-in mice are described (1). Cells were cultured in Dulbecco’s Modified Eagle Medium (Thermo Fisher, Cat. No. 12100046) containing 10% fetal bovine serum, penicillin (100 U/ml)/ streptomycin (100 mg/ml). MEF and RPE cells were transfected with Lipofectamine 3000 according to the manufacturer. Lentivirus particles were produced in 293T cells and infected in target cells as described previously (5). After lentiviral transduction, cells were sorted on SONY SH800Z cell sorter to collect the lowest expressing cells to avoid overexpression artefacts. In the case of GFP-TTBK2, viral transduced cells were selected using Puromycin selection. RILPL1-GFP was induced by 1μg/ml doxycycline 24h after transfection.

### Live imaging

MEF R1441C cells were plated on 8-well dishes at 3×10^4^ cells per each well (Nunc™ Lab-Tek™ II Chambered Coverglass from Thermo Fisher Scientific). Cells were cultured overnight with complete media including 10% FBS. To observe cilia formation, cells were washed 3 times and transferred to Leibovitz’s L-15 Medium, no phenol red (Thermo Fisher Scientific) with 200 nM MLi-2 or DMSO for 2hr before imaging. For monitoring cilia loss, cells were washed with PBS 3 times and cultured with complete media lacking serum for 20hr. Then cells were transferred to L-15 Medium with 10% FBS ± MLi-2; after 2hr culture, cells were imaged for 8hr. For MLi-2 washout, ciliated cells were washed and transferred to L-15 Medium ± MLi-2. Images were captured using Metamorph software at every 15 minutes in 37C (PeCon Tempcontrol 37-2) using confocal spinning disk system (Leica DMI 6000 B equipped with YOKOGAWA CSU-W1 Confocal Scanner Unit) with ANDOR iXon Life 897 EMCCD camera and 63x glycerol immersion objective (HC PL APO CS2 63x/1.30 GLYC, leica).

### Light microscopy

Cells were plated on glass coverslips and starved for 24 h. Cells were then fixed with cold methanol for 5 min, blocked with 1% BSA in PBS, and stained with indicated antibodies. Primary and secondary antibody incubations were for 1 h at RT. Highly cross-absorbed H+L secondary antibodies (Life Technologies) conjugated to Alexa Fluor 488, 568, or 647 were used at 1:5000. Nuclei were stained with 0.1 μg/ml DAPI (Sigma). Glass coverslips were mounted onto slides using mowiol. All images were obtained using Metamorph software with a spinning disk confocal microscope with 100× 1.4-NA oil-immersion objective at room temperature.

### Image analysis

For live cell imaging analysis, Z stacks of captured pictures were subject to maximum intensity projections and areas of interest were cropped by Fiji (National Institutes of Health). A decapitation event was designated if a vesicular SSTR3-GFP+ structure appeared near the tip of the cilium. Numbers and timing of cilia generation, loss or decapitation were counted manually. Cilium length was measured using Cell profiler (27). After highlighting cilia structures by enhancing neurite features, cilia were identified as objects. Cilia were skeletonized, then pixel lengths were calculated. For quantifying TTBK2 at the centrosome, CellProfiler software was used as follows. Maximum intensity projection images of CEP164 staining were used to identify mother centrioles. The mean intensities of TTBK2 staining were calculated within the circular areas around the M-centriole of 1.5 μm diameter and 10 μm diameter. The inner ring was used to determine TTBK2 local concentration whereas the outer ring provided information related to overall TTBK2 levels. The ratios of these two values were used to determine TTBK2 concentration at the M-centriole. EHD1 was analyzed by the same method except that gamma tubulin staining was used to determine centrosomal localization and thus each cell had two gamma tubbulin circles that were analyzed.

### Statistics

Graphs were made using GraphPad Prism 6 software. Error bars indicate SEM. A Student’s unpaired *t* test was used to test significance. Two-tailed P values <0.05 were considered statistically significant.

## Acknowledgements

This research was funded by grants to SRP from the U.S. National Institutes of Health (DK37332) and from the Michael J. Fox Foundation for Parkinson’s research.

---

**Supplemental Figure 1.**
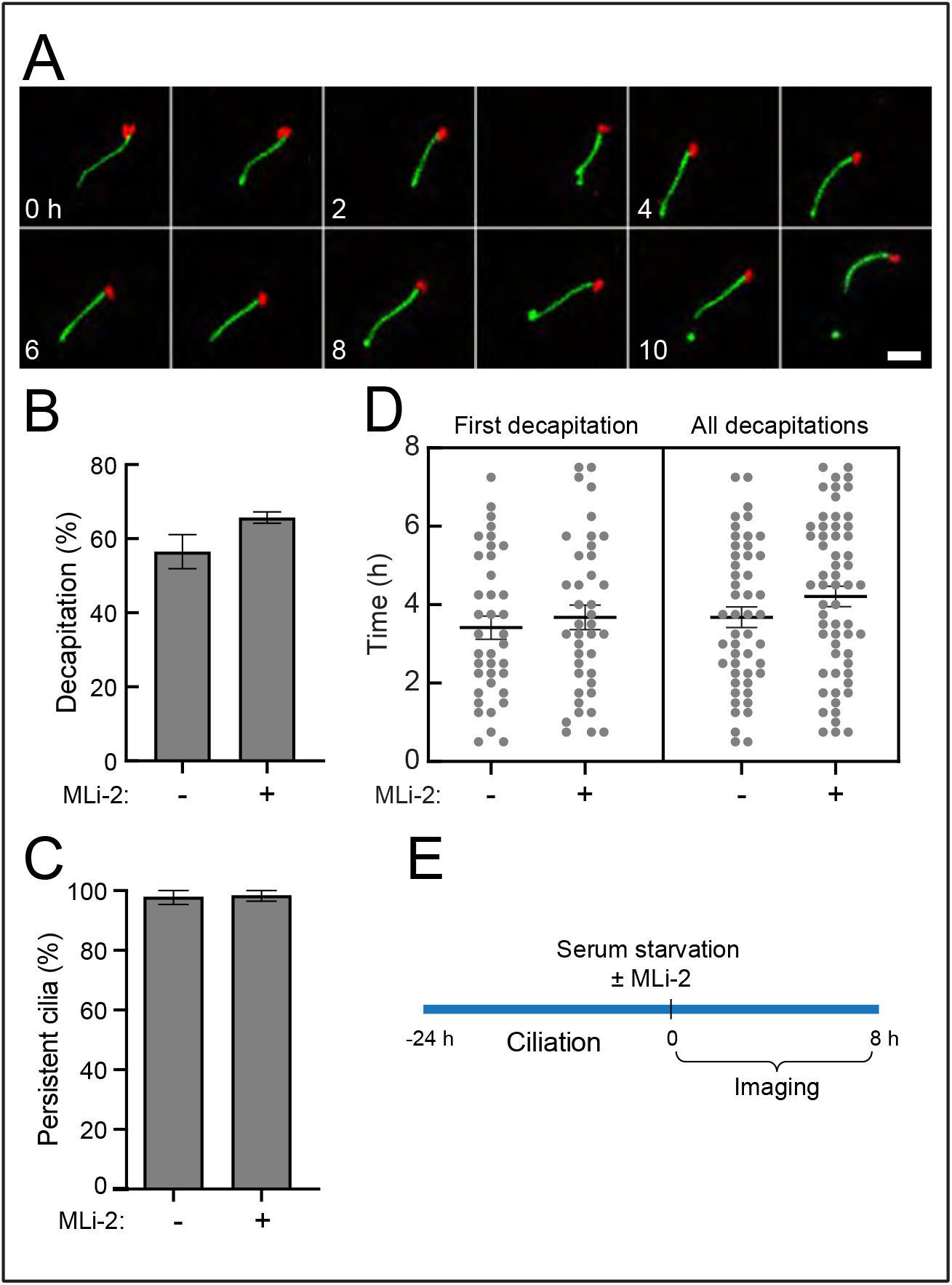
Cilia formed with or without pathogenic LRRK2 activity are stable in the absence of serum. **A.** Time-lapse images of R1441C LRRK2 MEF cells stably expressing SSTR3-GFP (green) and mKO2-PACT (red). Cells were serum starved ± MLi-2 for 24hr then imaged every 15 minutes. Scale bar, 5μm. **B., C.** Percentage of cilia showing any decapitation events (P=0.13) and probability of cilia persistence (P=0.86) during 7hr imaging ± MLi-2. Values represent the mean ± SEM from 3 independent experiments, each containing >19 cells. Significance was determined by t-test. **D.** Timing of first decapitation of each cilium (P=0.54) and all decapitation events (P=0.15) are shown. Values represent the mean ± SEM. Significance was determined by t-test. **E.** Timeline of this imaging experiment.

**Supplemental Figure 2.**
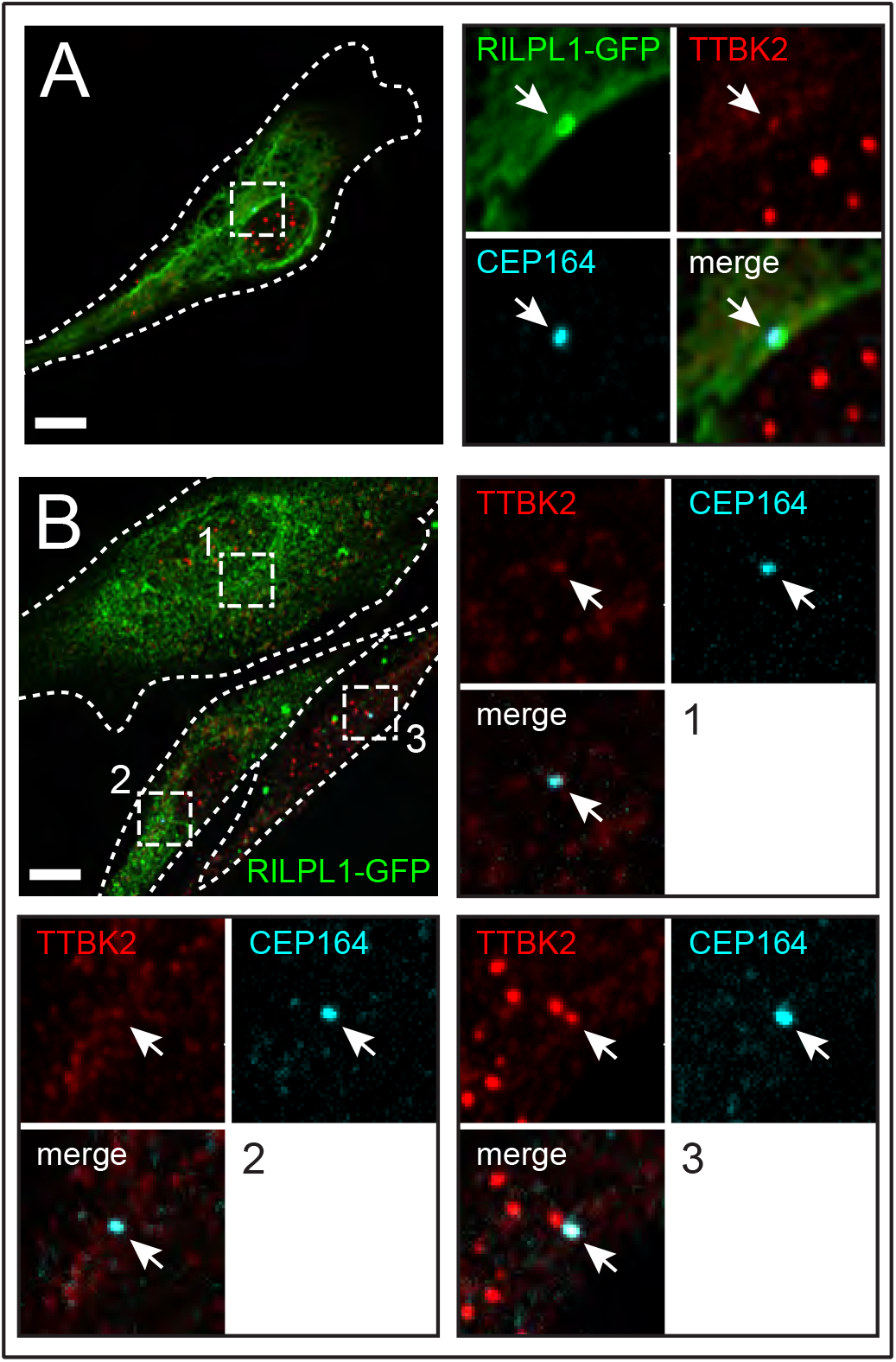
TTBK2 localization in serum starved RPE cells over-expressing RILPL1-GFP as in Figure 6D. Individual cells are outlined in A and B; enlarged regions are boxed. In B, #1 and #2 cells express more RILPL1-GFP than #3; the surrounding enlargements are labeled to indicate cell #1, 2 or 3. RILPL1-GFP was detected with chicken anti-GFP (green); CEP164 was detected using mouse anti-CEP164 (turquoise); rabbit anti-TTBK2 staining is shown in red. Scale bar, 10μm. Arrows indicate the location of CEP164 at the mother centriole. Note that centriolar RILPL1 staining is often lost in methanol fixed cells compared with paraformaldehyde fixation, thus merges in B are 2-color, to highlight TTBK2 and CEP164.

